## Supplementary material for "FMed-Diffusion Federated Learning on Medical Image Diffusion": source: main.tex

\documentclass[lettersize,journal]{IEEEtran}

\usepackage{amsmath,amsfonts}
\usepackage{algorithmic}
\usepackage{algorithm}
\usepackage{array}
\usepackage[caption=false,font=normalsize,labelfont=sf,textfont=sf]{subfig}
\usepackage{textcomp}
\usepackage{stfloats}
\usepackage{url}
\usepackage{verbatim}
\usepackage{graphicx}
\usepackage{cite}
\usepackage{graphicx}

% \usepackage{amsmath}
\newcommand\norm[1]{\left\lVert#1\right\rVert}
% \usepackage{mathtools}

\usepackage{subfig}

\usepackage{float}

% \usepackage{algorithm}
\usepackage[noend]{algpseudocode}
\hyphenation{op-tical net-works semi-conduc-tor IEEE-Xplore}
% updated with editorial comments 8/9/2021

\begin{document}
 
\title{FMed-Diffusion Federated Learning on Medical Image Diffusion}

\author{Murukessan~Perumal~,
        and M~Srinivas~
        % <-this % stops a space
\thanks{The authors would like to thank the Ministry of Education Government of India, NIT Warangal, and the NITW/CS/CSE-RSM/2018/908/3118 project for supporting this study.}
\thanks{(Corresponding author: Murukessan Perumal)  
% note need leading \protect in front of \\ to get a newline within \thanks as
% \\ is fragile and will error, could use \hfil\break instead.
}% <-this % stops a space
\thanks{Murukessan Perumal and Dr.M Srinivas are with the Department of Computer Science and Engineering at the National Institute of Technology at Warangal in India. \protect\\}}

% The paper headers
% \markboth{Please submit the manuscript to the Special Issue on Trustworthy Federated
% Learning}%
% {Shell \MakeLowercase{\textit{et al.}}:FMed-Diffusion Federated Learning on Medical Image Diffusion}

% \IEEEpubid{0000--0000/00\$00.00~\copyright~2021 IEEE}
% Remember, if you use this you must call \IEEEpubidadjcol in the second
% column for its text to clear the IEEEpubid mark.

\maketitle


\begin{abstract}
Medical data is not available for public access due to privacy concerns of the patients and the stakeholders' trustworthiness. However, Artificial Intelligence, especially all deep-learning models, is data-hungry and fails to produce clinically relevant results without much data. Moreover, augmentation strategies are deployed to overcome the less data hurdle. The promising future in this direction is generative AI-augmented data. The chat-GPT and DALLE-2 have become commercial products leveraging the generative AI in Natural Language Processing and Computer Vision. The diffusion models have started giving many promising results in the generative AI in computer vision. And in medical imaging, they can potentially create synthetic data to augment the scarce dataset. Diffusion models coupled with federated learning can create synthetic data on a large scale without the need to violate data privacy. This synthetic dataset could be used for further training of deep learning models without the issues of patients' identity theft from reverse engineering data. A Federated Learning paradigm of diffusion models has been proposed to overcome this hurdle and its related challenges. Our work focuses on diffusion models in federated learning settings. We named our novel model FMed-Diffusion or Federated Learning on Medical Image Diffusion. We trained our model under distributed federated settings imitating real-world clinical settings. We have achieved impressive results over three medical image datasets, APTOS 2019 Blindness Detection, Retinal OCT Detection, and COVID-CT Detection in the federated setting on par with the traditional training. Our model FMed-Diffusion has achieved an FID score of 7.1821 on the APTOS 2019 Blindness Detection dataset, an FID score of 8.8154 on the Retinal OCT Detection dataset, and an FID score of 7.4486 on the COVID-CT Detection dataset.
\end{abstract}

\begin{IEEEkeywords}
Federated Learning, Diffusion models, Medical Imaging, Distributed Learning, Generative-AI.
\end{IEEEkeywords}

\section{Introduction}
Federated learning has emerged as a viable alternative to privacy concerns in the healthcare sector. It deploys federated distributed learning through the global aggregation of model weights and local training with the aggregated model weights. This strategy avoids sharing patients' critical data, which may violate their privacy concerns. The gadget ecosystem that has exploded through the Internet of Things(IoT) in medical devices like smartwatches is one example that tracks medical data and monitors patients' health. In such a scenario, patient information privacy becomes critical due to the sensitive personal information involved. There are efforts to ensure the security of patients' data in such environments, for example, querying medical health records in multi-user settings \cite{hatamizadeh2023gradient,10068787,li2022auditing}. Federated Learning solves the data privacy issue by sharing the aggregated global model with all the clients of different medical institutes participating in the learning instead of sharing the sensitive patients' data. Therefore, this distributed training eases the legal issues and complications of sharing medical records.

It all started with a seminal paper \cite{mcmahan2017communication} that termed their distributed way of training to address learning from non-centralized data that forms most of the real-world scenario of privacy-sensitive data as Federated Learning. There is local training and aggregation of the locally trained models into a global model in the federated settings, and the cycle repeats until convergence is achieved satisfactorily. There are various strategies involved in the aggregation process in global parameter sharing. And there are strategies involved in avoiding the local minima during the local training with the limited data per client institution. There are common challenges involved in the federated learning setup. And these challenges reflect most of the real-world scenarios and preclude the custom data processing possible in a centralized data center. This makes the federated learned models work on most real-time scenarios. This is more pertinent to the medical domain, where patient privacy concerns force such a situation. This \cite{nguyen2022federated} survey gives a comprehensive overview of Federated Learning in Medical IoT devices for smart healthcare.

There are works \cite{fang2022robust,gao2022feddc,huang2022learn,jain2023federated,li2022federated,mendieta2022local,michieli2021all,qu2022rethinking,zhang2021federated,hosseini2023proportionally,li2021model} addressing the various issues and challenges involving federated learning. They address the issues of non-IID data distribution, catastrophic forgetting due to local training, heterogeneous client models, commercial setup of client devices, fairness among the clients, architectural choices in clients, the disparity in feature representations in clients, local drift in client updates, and permutation invariance of neural networks. Providing solutions at the client or server level differs from work to work. Previously, Google claimed the replacement of radiologists with deep learning models, but their model developed in the lab failed miserably in real-time. Most deep learning models produced in labs for unstructured medical modalities such as X-Rays, CTs, Medical Health Records, Optical Coherence Tomography (OCT), Fundus images, and so on do not do well in real-time scenarios. They are not deployable in real-time clinical environments. This failure is mainly part of not reproducing the actual clinical settings in the research labs. However, Federated Learning could be a blessing in disguise to enforce such clinical constraints in the research labs, especially in medical imaging \cite{yan2023label,diaz2023study,rieke2020future}. Even though there are recent works in Federated Learning in the medical imaging domain, the exploration of the same is not done for the latest diffusion models. We try to fulfill the gap with our  Federated Learning on Medical Image Diffusion (FMed-Diffusion) model.

In our work, we have trained the novel FMed-Diffusion on Diabetic Retinopathy (Blindness-Detection) \cite{KaggleAPTOS}, OCT on retinal scans (OCT-Scan) \cite{kermany2018labeled}, and COVID-CT scans (COVID-CT) \cite{soares2020sars} datasets. We have achieved realistic-looking synthetic images by training in the federated setting and on par with images by training in centralized data center settings. We compare and contrast federated and centralized learning on those datasets and diffusion models. We have replicated the exact Federated Learning settings with server-client architecture and as distributed training. The diffusion models are the latest set of training strategies that are used for generative imaging. It is radically different in its methodology from previous state-of-the-art generative models such as Generative Adversarial Networks(GAN) \cite{goodfellow2020generative}, Flow-based methods \cite{rezende2015variational}, and Variational Auto-Encoders (VAE). The diffusion models came to the limelight through the groundbreaking work \cite{ho2020denoising}.

Further, works \cite{croitoru2023diffusion,dhariwal2021diffusion,saharia2022photorealistic,rombach2022high,qin2023class,hu2023self} consolidated and corroborated the status of the diffusion models as the latest trending models in the field of computer vision and generative AI. There are applications of diffusion models in the medical imaging domain \cite{kazerouni2022diffusion}. Their work is a comprehensive overview of various applications of different diffusion models in the medical imaging domain. Particularly in dermatology, where the deep learning models have achieved human expert level artificial expertise, there is the application of diffusion model \cite{akrout2023diffusion} for next-generation data augmentation through latent diffusion. They control the synthetic image generation through prompt text that aligns well with the generated image. A model trained solely on this synthetic dataset achieves classification accuracy in par with the one trained on real data.  

The pioneer diffusion model, named Denoising Diffusion Probabilistic model \cite{ho2020denoising}, has a time-scaled forward diffusion phase and a denoising or progressive error-removing reverse diffusion stage. The number of time steps involved in the forward and backward stage add a computational burden on the training process, much like the sequential training process of RNN or LSTM. However, diffusion models have proven to be suitable for generative imaging with no mode collapse or unstable training that degrades the GAN models and the constraints on models, such as in the flow-based methods. And they are better at distinguishing boundary information than their flow-based counterparts at the cost of computation time and sampling efficiency. Our proposed model FMed-Diffusion is a denoising diffusion probabilistic model trained in federated learning settings that has achieved impressive Frechet Inception Distance (FID) scores on three standard datasets. To our knowledge, Diffusion models are still not explored in the space of Federated Learning in the literature. 

\section{FMed-Diffusion}
The proposed novel FMed-Diffusion is a Denoising Diffusion Probabilistic model trained on medical imaging datasets in a Federated Learning approach. The Federated Learning setup is shown in Figure \ref{fig:pm}. The federated setup has a model aggregating server that collects locally trained model weights from the clients and does a federated average of those weights using the FedAvg strategy. There are about three clients, and all do localized training of the denoising diffusion probabilistic models in their datasets. The original datasets are divided to replicate the non-IID(Independent and Identically Distributed) nature of the real-time multiple clinical settings and their privacy concerns. The server is provided only with the model weights from the respective clients, whereas the data used for training in each client is unknown to the server and other clients. There is a single server and three clients in our federated settings.
% trim={left bottom right top}
\begin{figure}
    \centering
    \includegraphics[width=\columnwidth,trim={2cm 12cm 3cm 0},clip]{FL1.drawio.pdf}
    \caption{Federated Learning: Distributed Network Setup}
    \label{fig:pm}
\end{figure}

Each client does the localized training on the medical imaging datasets. It learns to reduce the loss locally and then communicates the weights of the local model to the server. The server collects all the weights from the clients and aggregates them through FedAvg algorithm, which averages the weights and provides the global average weights to all the clients. The clients again repeat the process for every round. Due to the diffusion models' complexity, we experimented with three clients and one FedAvg server. The model used in the denoising process to predict the noise to be removed over timesteps is the denoising diffusion probabilistic model, U-Net. It predicts a noise in equal dimensions to the image at every time step to be denoised from the latent noise progressively over time steps.

Figure \ref{fig:pdm}. shows the denoising steps on sample images from Blindness-Detection and OCT-Scan datasets. As shown in the figure, the forward pass consists of progressively adding noise until the images become pure noise. Then a U-Net model is trained to recover the original image over the timesteps by predicting the noise to be removed from the image, given the image+noise and the timestep. The U-Net is trained over timesteps to predict the image from pure latent noise. This is an end-to-end deep learning with the overhead number of timesteps per forward and backward diffusion stages. The forward pass here refers to the progress over each timestep in adding noise, and the backward pass refers to recovering the image from the noise.  
% trim={left bottom right top}
\begin{figure*}
    \centering
    \includegraphics[width=2.0\columnwidth,trim={0cm 5cm 4cm 0cm},clip]{Diff.pdf}
    \caption{FMed-Diffusion: Overview of proposed model Federated Learning on Medical Image Diffusion}
    \label{fig:pdm}
\end{figure*}

\begin{equation}
  % \scalebox{0.8}{
    Prob(img_t|img_{t-1}) := \mathcal{N}(img_t;\sqrt{1-V_t}img_{t-1},V_tI)
    % }  
    \label{eq:noise}
\end{equation}

% \begin{equation}
  % \scalebox{0.8}{
  \begin{gather}
        \alpha_t := 1-V_t  \\
        \overline{\alpha_t} := \prod^t_{s=1} \alpha_t \\
        Prob(img_t|img_0) := \mathcal{N}(img_t;\sqrt{\overline{\alpha_t}}img_0,(1-\overline{\alpha_t})I)
        \label{eq:repnoise}
  \end{gather}
    % }
    % \label{eq:noise}
% \end{equation}

The noise added to the image progressively over the timesteps in the forward pass of the diffusion model training follows Equation \ref{eq:noise}, where the probability distribution $Prob$ of a noisy image to transit from $t-1$ step to $t$ follows the Multivariate Gaussian Distribution with variance $V_t$ for time step $t$. And a scaling reduction of the mean is taken as $img_{t-1}$ by a factor of $\sqrt{1-V_t}$. The overall noisy sampling progresses towards pure latent noise by scaling the mean and perturbing the image with the noise of variance $V_t$. Since this is a multivariate distribution, there is a covariance matrix in place of variance, given by $V_tI$ at time step $t$. The covariant matrix is diagonal and is scaled variance at time step $t$ over identity matrix $I$. This is to have an equal variance for all latent space dimensions. Even though this is a simple equation, calculating it over multiple steps becomes tedious and cumbersome even as computations. Therefore reparameterization trick is used to arrive at Equation \ref{eq:repnoise}. From Equation 4, one can easily estimate the noisy image at $t$ timestep directly without going through from the start to $t$ time step. The process of forward diffusion is necessary to create targets for the reverse diffusion process. The diffusion process is a Markov chain with the weak assumption that each timestep depends only on its previous timestep. The assumption of the Markov chain makes Equation 4 valid to compute the noise at the $t$ timestep at a time.

% % \begin{equation}
%   % \scalebox{0.8}{
%   \begin{gather}
%         \alpha_t := 1-V_t  \\
%         \overline{\alpha_t} := \prod^t_{s=1} \alpha_s \\
%         Prob(img_t|img_0) := \mathcal{N}(img_t;\sqrt{\overline{\alpha_t}}img_0,(1-\overline{\alpha_t})I)
%         \label{eq:repnoise}
%   \end{gather}
%     % }
%     % \label{eq:noise}
% % \end{equation}

The reverse diffusion process is to learn the noise given the image and the timestep. A U-Net model is used since the reverse diffusion process step needs to predict an error equal to the dimension of the image. The U-Net model is an encoder-decoder model that makes an image-to-image prediction. It is very well established in the field of medical image segmentation, where each pixel in the image needs to be assigned a label. It employs skip connections to enrich the prediction with less lossy downscaling and effective upscaling in the U-shaped path. Deploying the diffusion model U-Net in a federated learning environment is a challenge in itself. The Federated setting has challenges to address, such as distributed training, communication between a server and its clients, and the nature of data distributed among the clients.

\section{Results and Discussion}
% \centering
\begin{figure*}
    \centering
    % % trim={left bottom right top}
    % \includegraphics[width=2.0\columnwidth,trim={0cm 0cm 0cm 0cm},clip]{Dr24.png}

    % \centering
    % \hspace{1cm}
    \subfloat[]{		         
    \includegraphics[width=0.75\textwidth,height=6cm,trim={0cm 0cm 0cm 0cm}]{Dr24.png}
    } \\
    % \newline
    % \hfill
    % \centering
    % \hspace{2cm}
    % \centering
    \subfloat[]{
    \includegraphics[width=0.75\textwidth,height=6cm,trim={0cm 0cm 0cm 0cm}]{OCT24.png}
    } \\
    % \newline
    % \hfill
    % \centering
    \subfloat[]{		
    \includegraphics[width=0.75\textwidth,height=6cm,trim={0cm 0cm 0cm 0cm}]{CT24.png}
    }
    
    \caption{Generated sample images from FMed-Diffusion trained on (a) is Blindness Detection, (b) is OCT-Scan, and (c) is COVID-CT datasets}
    \label{fig:genimages}
\end{figure*}

\begin{figure}
    % \centering
    % % trim={left bottom right top}
    % \includegraphics[width=2.0\columnwidth,trim={0cm 0cm 0cm 0cm},clip]{Dr24.png}

    \centering
    \subfloat[]{	    \includegraphics[width=0.45\columnwidth,height=3cm]{C_s16.png}
    }
    % \newline
    \hfill
    \subfloat[]{		\includegraphics[width=0.45\columnwidth,height=3cm]{F_s46.png}}
    % \newline
    % \hfill
    \\
    \subfloat[]{		\includegraphics[width=0.45\columnwidth,height=3cm]{C_s12.png}}
    \hfill
    \subfloat[]{	    \includegraphics[width=0.45\columnwidth,height=3cm]{F_s11.png}
    }
    \\
    % \newline
    % \hfill
    \subfloat[]{		\includegraphics[width=0.45\columnwidth,height=3cm]{C_s16_COVID.png}}
    % \newline
    \hfill
    \subfloat[]{		\includegraphics[width=0.45\columnwidth,height=3cm]{F_s187.png}}
    
    \caption{Comparison of images generated by centralized training of diffusion (a,c,e) vs. FMed-Diffusion (b,d,f)}
    \label{fig:comp}
\end{figure}

The experiments were conducted on a DGX-A100 workstation. The Federated Learning(FL) was simulated using the Flower \cite{beutel2020flower} Python package. The experimental setup is very close to the real-time distributed environment in medical organizations. The Hugginface Diffusers library \cite{von-platen-etal-2022-diffusers} was used for Diffusion model training and the generation of samples. The Federated Learning server used FedAvg algorithm to aggregate the global model from all the client models. We fixed the number of clients to three due to hardware limitations and the time required for simulation in diffusion models. Each client used a slice of the GPU and therefore is separate from each other and the server in the process, resembling the real-time distributed scenario closely. The dataset is split into three parts and is spread across each client.

\begin{figure*}
    % \centering
    % % trim={left bottom right top}
    % \includegraphics[width=2.0\columnwidth,trim={0cm 0cm 0cm 0cm},clip]{Dr24.png}

    \centering
    \subfloat[]{		         
    \includegraphics[width=0.3\textwidth,height=4cm,trim={1cm 1cm 1cm 1cm},clip]{C1TrainLoss.jpg}
    }
    % \newline
    % \hfill
    \subfloat[]{		\includegraphics[width=0.3\textwidth,height=4cm,trim={1cm 1cm 1cm 1cm},clip]{C2TrainLoss.jpg}}
    % \newline
    % \hfill
    \subfloat[]{		\includegraphics[width=0.3\textwidth,height=4cm,trim={1cm 1cm 1cm 1cm},clip]{C3TrainLoss.jpg}}
    
    \caption{Training loss per client for the first ten rounds(with generated names and colors) logged with Wandb \cite{wandb} at client1 (a), client2 (b), and client3 (c)}
    \label{fig:trainData}
\end{figure*}

To resemble the real-world scenario of non-IID data, we separated the datasets into three parts, breaking the classification labels' IID assumption. The number of samples in each client was also kept different to resemble the read-world distributed diffusion training process. The training hyperparameters for the clients are image size, batch size, number of epochs in each client, learning rate, and number of rounds for the server. The mixed precision was used to speed up training further. The initial learning rate and the number of diffusion time steps were fixed as 0.0001 and 1000 for all experiments. The image size, batch size, epochs, and rounds for OCT-Scan and COVID-CT datasets are 224, 22, 5, and 10, respectively. The same for the Blindness-Detection dataset differs to 512, 6, 10, and 20. Due to the higher resolution of Fundus images, their training parameters are different. 

The Blindness Detection dataset from Kaggle \cite{KaggleAPTOS} is used to detect Diabetic Retinopathy (DR), a significant cause of blindness due to diabetes, from the fundus images. The fundus images are graded from 0-4 for the severity of Diabetic Retinopathy, indicating 0-no DR, 1-Mild DR, 2-Moderate DR, 3-Severe DR, and 4-Proliferative DR. The number of training images in the classes 0, 1, 2, 3, 4 are 1805, 999, 370, 295, 193 respectively. There are a total of 3662 train images and 1928 test images. The test images have no labels. In the FL experiment on this dataset, client one has 1805 images of class 0, client two has 999 images of class 2, and client three has 858 images of class 1,3,4. This distribution of data is a non-IID and also imbalanced replicating real-world scenarios. 

The OCT-Scan dataset \cite{kermany2018labeled}, a Mendeley dataset of Retinal Optical Coherence Tomography (OCT) images that capture high-resolution cross sections of the retinas of living human patients from Shiley Eye Institute of the University of California San Diego, the California Retinal Research Foundation, Medical Center Ophthalmology Associates, the Shanghai First People’s Hospital, and Beijing Tongren Eye Center between July 1, 2013, and March 1, 2017. The images are classified into abnormalities such as Choroidal NeoVascularization (CNV), Diabetic Macular Edema (DME), Multiple Drusen, and Normal retina. There are 83484 train images, of which 37205 are CNV, 11348 are DME, 8616 are Drusen, and 26315 are Normal. There are 968 testing images, with 242 images from each class. In the FL experiment on this dataset, client one has all 37205 CNV images, client two has all 26315 Normal images, and client three has 8616 combining DME and Drusen images. Thus the FL setup resembles the real-world setting of distributed, non-IID, and imbalanced data.

The COVID-CT dataset \cite{soares2020sars} has 2D slices of chest CT scans of actual patients for SARS-CoV-2 identification taken from patients in Sao Paulo - Brazil. The dataset is divided into three classes COVID, Healthy, and Others. There are about 2168 COVID images, 758 Healthy images, and 1247 Other findings. There is no train-test split in the dataset. In our FL experiment on this dataset, client one has all COVID images, client two has Healthy images, and Client three has all Other finding images. This is by the actual clinical settings where different clinics may have different types and numbers of samples of medical images.

The experiments were conducted with three clients with the above data splits and a server to facilitate aggregating, averaging, and distributing the global model weights from all the clients' weights. The number of total epochs can be a rough estimate of the number of local epochs at each client times the number of rounds in the server. The algorithm \ref{algo:FL} explains the Federated Learning of Denoising Diffusion U-Net model in a detailed way. A single server and three clients are in a distributed environment employing federated learning. Initially, the server sends the global model to all three clients. The clients run as separate processes, receive the global model weights, and start learning from their local data with the U-Net model's initial weights as the global model weights. In the local training process, each image from the local private dataset is used as a starting condition to generate a noise-added image following Equation \ref{eq:repnoise}. It is repeated for $T$ timesteps by following a uniform distribution on timestep. The U-Net is trained to predict the noise given the noisy image and the timestep of the induced noise. The U-Net error prediction is optimized to equal the added error by the forward diffusion process. The U-Net takes the noisy image and predicts the noise latent of equal dimensions. After this local learning for several epochs in each client, the newly learned weights of each client are sent to the server for averaging by the FedAvg algorithm. The averaged global weights are again shared with the clients by the server. And this cycle is repeated for several rounds. Finally, after a sufficient number of server rounds and, in each round, local training by each client, the training is stopped. The images are then sampled from the trained model by denoising them over several timesteps from the initial latent noise to generate high-quality images. The sampling step involves reducing the error predicted by U-Net at each timestep after rescaling them to get the mean and adding it with the unit normal distribution scaled by the standard deviation of the reverse diffusion at each timestep. This process gives rise to a lot of variation in sampling and enriching samples of high diversity and quality.

The sample images generated using the model trained in FL settings on the three datasets are shown in Figure \ref{fig:genimages}. Figure \ref{fig:genimages} (a) shows a sample of all the synthetic images generated by the proposed diffusion model FMed-Diffusion trained on the Blindness Detection dataset. It can be seen by first-hand manual inspection that they look very realistic and have a lot of variation among themselves. We also corroborated our qualitative human assessment with a quantitative measure called Frechet-Inception-Distance (FID) \cite{NIPS2017_8a1d6947,Nicki_Skafte_Detlefsen_and_Jiri_Borovec_and_Justus_Schock_and_Ananya_Harsh_and_Teddy_Koker_and_Luca_Di_Liello_and_Daniel_Stancl_and_Changsheng_Quan_and_Maxim_Grechkin_and_William_Falcon_TorchMetrics_-_Measuring_2022} given by the Equation \ref{eq:FID}. Figure \ref{fig:genimages} (b) shows a collection of sampled images from the proposed model FMed-Diffusion trained on the OCT-Scan dataset. The collection of OCT images looks very realistic on manual examination. The results are consolidated with FID for a quantitative evaluation. Figure \ref{fig:genimages} (c) shows the sample of generated images from the proposed model FMed-Diffusion trained on the COVID-CT dataset. The images look promising and of higher quality from the initial visual inspection.
Further, these images are consolidated with the FID score to measure image generation by diffusion quantitatively. Figure \ref{fig:comp} compares the images generated using centralized settings and federated settings. The left column of images is generated using centralized settings where the training diffusion model has access to the whole dataset. At the same time, the right column of images is generated by FMed-Diffusion in a federated setting. On visual inspection, it can be observed that the proposed model FMed-Diffusion has generated samples of high quality on par with centralized training of the same. Also, the proposed model FMed-Diffusion achieved impressive FID scores on the three datasets Blindness Detection, OCT-Scan, and COVID-CT as 7.1821, 8.8154, and 7.4486, respectively. The details are tabulated and plotted as a line graph for visualization in Table \ref{tab:FID} and Figure \ref{fig:FID}, respectively. 

\begin{equation}
    FID(P,Q) = \sqrt{\norm{\mu_p - \mu_q}_2^2 + Tr(\mathcal{E}_p+\mathcal{E}_q-2\sqrt{(\mathcal{E}_p\mathcal{E}_q))}}
    \label{eq:FID}
\end{equation}

The FID score is calculated by the Equation \ref{eq:FID}, where the distance between two Multivariate Gaussian distributions is calculated. Here P and Q are the two Multivariate Gaussian distributions, and $\mu_p$ and $\mu_q$ are their respective means, $\mathcal{E}_p$ and $\mathcal{E}_q$ are their respective covariance matrices, and $Tr$ is the trace of a matrix. It essentially captures the distance between the mean and the variance in a multivariate setting. The distribution variables come from the features extracted through the Inception model trained on Imagenet, hence the name. The features are used as multivariate continuous random variables for the distributions. FID calculates the distance between the actual data distribution and that of the synthetically generated by FMed-Diffusion. The FMed-Diffusion sampled the images and generated samples in equal numbers to the test datasets in the Blindness Detection and the OCT-Scan datasets. In contrast, generated samples were equal to the COVID-CT dataset's training dataset; no train-test split was given. The sampled images are then used to calculate the FID Score with the actual test datasets for all datasets except COVID-CT, where the training dataset was used.

Table \ref{tab:FID}. shows the FID Score achieved by FMed-Diffusion on all three datasets. A lower FID Score indicates that the model has performed well on sampling and has matched more with the test datasets. Our proposed model is on par after a rough comparison with the FID scores achieved by generative models in the medical imaging \cite{moghadam2023morphology,horvath2022metgan,Hou:23}. Visual representation of the FID scores as a line graph in Figure \ref{fig:FID}. shows the model has performed better on Blindness detection than on COVID-CT and the OCT-Scan datasets. This could be attributed to the higher resolution of the Fundus images compared to other datasets and the noisy images in the OCT-Scan training dataset. This comparison is rough and weak since the number of generated samples differs in each dataset. The training loss per client for the first ten rounds is shown in Figure \ref{fig:trainData}. All the clients try to learn from the local data by minimizing the loss, that is, minimizing the distance between the predicted noise by U-Net and the target noise by a forward diffusion process. Wandb \cite{wandb} pulled the data in Figure \ref{fig:trainData} from the FMed-Diffusion trained over the Blindness Detection dataset for the first ten rounds. It shows the loss reduction over the number of steps in the epochs.

\begin{table}
\centering
  \caption{Frechet Inception Score (FID) achieved by FMed-Diffusion on all three datasets}
  \begin{tabular}{|c|c|}
      \hline
      \textbf{Dataset} & \textbf{FID Score by FMed-Diffusion} \\
      \hline
      Blindness Detection & 7.1821 \\
      \hline
      OCT-Scan & 8.8154 \\
      \hline
      COVID-CT & 7.4486 \\
      \hline
  \end{tabular}
  \label{tab:FID}
\end{table}

\begin{figure}
    \centering
    \includegraphics[width=\columnwidth]{line_graph_FID.png}
    \caption{Line Plot of FID Scores by FMed-Diffusion on all three datasets}
    \label{fig:FID}
\end{figure}


\begin{algorithm}
\caption{FMed-Diffusion}
\begin{algorithmic}[1]
\State \textbf{Server:}
\State\hspace{1em} Initialize global model $M_0$
\State\hspace{1em} Send $M_0$ to all clients
\\
\State \textbf{Client $i$ ($i = 1, 2, 3$):}
\State\hspace{1em} Receive global model $M_0$
\State\hspace{1em} Initialize local model $L_i \gets M_0$
\State\hspace{1em} Sample local data $D_i$
\State\hspace{1em} Train $L_i$ on $D_i$ for $N$ epochs
\State\hspace{2em}  \textbf{For each Epoch in $N$}
\State\hspace{3em}  Diffusion Learning with $T$ timesteps
\State\hspace{3em}  \textbf{For $img_0$ in $D_i$}
\State\hspace{4em}  \textbf{repeat} $T$
\State\hspace{5em} $t \sim UniDist({1-T})$
\State\hspace{5em} $\epsilon \sim \mathcal{N}(0,I)$
\State\hspace{5em} $\epsilon_\theta \hookrightarrow UNetPrediction$
\State\hspace{5em} Optimise $\theta$ with gradient
\State\hspace{5em} $\nabla_\theta \norm{\epsilon - \epsilon_\theta(\sqrt{\overline{\alpha_t}}img_0 + \sqrt{1-\overline{\alpha_t}}\epsilon, t)}^2$
\State\hspace{1em} $L_i$ learns from the local loss function
\\
\State \textbf{Client $i$:}
\State\hspace{1em} Send $L_i$ to the server
\\
\State \textbf{Server:}
\State\hspace{1em} Receive $L_i$ from client $i$
\State\hspace{1em} Aggregate client models: 
\State\hspace{1em} $M_{\text{new}} \gets \frac{1}{3} \cdot (L_1 + L_2 + L_3)$
\State\hspace{1em} Update global model: $M_0 \gets M_{\text{new}}$
\State\hspace{1em} Send $M_0$ to all clients
\\
\State \textbf{Repeat above for $R$ no of rounds}
\\
\State \textbf{Sampling at every Client}
\State\hspace{1em} Starting from pure latent noise
\State\hspace{1em} \#Note $img_{1-T}$ are latent of dimension as $img_0$
\State\hspace{1em} $img_T \sim \mathcal{N}(0,I)$
\State\hspace{1em} \textbf{for $t$ = $T \hookrightarrow 1$}
\State\hspace{2em} $z \sim \mathcal{N}(0,I)$ if $t$!=1 else 0
\State\hspace{2em} \scalebox{1}{$img_{t-1} = \frac{1}{\sqrt{\alpha_t}}(img_t - \frac{1-\alpha_t}{\sqrt{1-\overline{\alpha_t}}}\epsilon_\theta(img_t,t))+\sigma_t z$}
\State\hspace{1em} New sample $\hookrightarrow img_0$
\end{algorithmic}
\label{algo:FL}
\end{algorithm}


\section{Conclusion}
We have obtained significant results on the diffusion models on medical imaging data in federated learning settings. And we applied distributed real-time approach to train those models. The models are evaluated with Freshcet Inception Distance which is used for qualitatively evaluating generative imaging models. The synthetic images sampled from the novel FMed-Diffusion exhibit high quality in manual visual inspection. The models are trained on three established medical imaging datasets. However, attacks on privacy, like reverse engineering data to patient information with any additional information along with the model weights sent to the server, need to be studied further. These generative models can be used for augmentations required to increase the synthetic dataset, which is pertinent to the data scarcity in the medical domain where the acquisition and collaboration of real-time quality data is a daydream that may not be practical. Further, we would like to work more on other variants of diffusion models, like latent diffusion in the federated learning paradigm, in the future. We would also like to explore the use cases of this generative modeling in augmenting datasets with synthetic images, which can be achieved through training in real-time federated distributed settings.

\bibliographystyle{ieeetr}
\bibliography{refs}

% biography section
% 
% If you have an EPS/PDF photo (graphicx package needed) extra braces are
% needed around the contents of the optional argument to biography to prevent
% the LaTeX parser from getting confused when it sees the complicated
% \includegraphics command within an optional argument. (You could create
% your own custom macro containing the \includegraphics command to make things
% simpler here.)
%\begin{IEEEbiography}[{\includegraphics[width=1in,height=1.25in,clip,keepaspectratio]{mshell}}]{Michael Shell}
% or if you just want to reserve a space for a photo:

\newpage

\begin{IEEEbiography}[{\includegraphics[width=1in,height=1.25in,clip,keepaspectratio]{muru.jpg}}]{Murukessan Perumal}
is a Research Scholar currently pursuing a Full-Time Ph.D. in the Department of Computer Science and Engineering at the National Institute of Technology, Warangal in India. He has obtained his bachelor's and master's in Computer Science Engineering from Anna University(CEG campus), Chennai, India. His research interest includes Medical Imaging, Machine Learning, Computer Vision, and Artificial Intelligence. 
\end{IEEEbiography}

% if you will not have a photo at all:
\begin{IEEEbiographynophoto}{M Srinivas}
is an Assistant Professor in the Department of Computer Science and Engineering at the National Institute of Technology, Warangal, in India. He obtained his Ph.D. from the Indian Institute of Technology (IIT), Hyderabad, India. He did post-doctorate research at ETS, Canada, and Academia Sinica, Taiwan. His research interests are Machine Learning, Medical Imaging, Computer Vision, Deep Learning, and Sparse Representation.
\end{IEEEbiographynophoto}
\vfill
\end{document}
