## Supplementary material for "FMed-Diffusion Federated Learning on Medical Image Diffusion": source: refs.bib

@ARTICLE{10068787,
  author={Xu, Chang and Chan, Zijian and Zhu, Liehuang and Zhang, Can and Lu, Rongxing and Guan, Yunguo},
  journal={IEEE Transactions on Sustainable Computing}, 
  title={EPPFM: Efficient and Privacy-Preserving Querying of Electronic Medical Records with Forward Privacy in Multiuser Setting}, 
  year={2023},
  volume={},
  number={},
  pages={1-12},
  doi={10.1109/TSUSC.2023.3257223}}

@inproceedings{mcmahan2017communication,
  title={Communication-efficient learning of deep networks from decentralized data},
  author={McMahan, Brendan and Moore, Eider and Ramage, Daniel and Hampson, Seth and y Arcas, Blaise Aguera},
  booktitle={Artificial intelligence and statistics},
  pages={1273--1282},
  year={2017},
  organization={PMLR}
}

@misc{KaggleAPTOS, title={APTOS 2019 Blindness Detection}, url={https://www.kaggle.com/competitions/aptos2019-blindness-detection}, author={Kaggle}, note = {Accessed: (05-15-2023)}} 

@article{kermany2018labeled,
  title={Labeled optical coherence tomography (oct) and chest x-ray images for classification},
  author={Kermany, Daniel and Zhang, Kang and Goldbaum, Michael and others},
  journal={Mendeley data},
  volume={2},
  number={2},
  pages={651},
  year={2018}
}

@article{soares2020sars,
  title={SARS-CoV-2 CT-scan dataset: A large dataset of real patients CT scans for SARS-CoV-2 identification},
  author={Soares, Eduardo and Angelov, Plamen and Biaso, Sarah and Froes, Michele Higa and Abe, Daniel Kanda},
  journal={MedRxiv},
  pages={2020--04},
  year={2020},
  publisher={Cold Spring Harbor Laboratory Press}
}

@article{yan2023label,
  title={Label-efficient self-supervised federated learning for tackling data heterogeneity in medical imaging},
  author={Yan, Rui and Qu, Liangqiong and Wei, Qingyue and Huang, Shih-Cheng and Shen, Liyue and Rubin, Daniel and Xing, Lei and Zhou, Yuyin},
  journal={IEEE Transactions on Medical Imaging},
  year={2023},
  publisher={IEEE}
}

@article{diaz2023study,
  title={Study of the performance and scalability of federated learning for medical imaging with intermittent clients},
  author={D{\'\i}az, Judith S{\'a}inz-Pardo and Garc{\'\i}a, {\'A}lvaro L{\'o}pez},
  journal={Neurocomputing},
  volume={518},
  pages={142--154},
  year={2023},
  publisher={Elsevier}
}

@article{ho2020denoising,
  title={Denoising diffusion probabilistic models},
  author={Ho, Jonathan and Jain, Ajay and Abbeel, Pieter},
  journal={Advances in Neural Information Processing Systems},
  volume={33},
  pages={6840--6851},
  year={2020}
}

@article{croitoru2023diffusion,
  title={Diffusion models in vision: A survey},
  author={Croitoru, Florinel-Alin and Hondru, Vlad and Ionescu, Radu Tudor and Shah, Mubarak},
  journal={IEEE Transactions on Pattern Analysis and Machine Intelligence},
  year={2023},
  publisher={IEEE}
}

@article{dhariwal2021diffusion,
  title={Diffusion models beat gans on image synthesis},
  author={Dhariwal, Prafulla and Nichol, Alexander},
  journal={Advances in Neural Information Processing Systems},
  volume={34},
  pages={8780--8794},
  year={2021}
}

@article{saharia2022photorealistic,
  title={Photorealistic text-to-image diffusion models with deep language understanding},
  author={Saharia, Chitwan and Chan, William and Saxena, Saurabh and Li, Lala and Whang, Jay and Denton, Emily L and Ghasemipour, Kamyar and Gontijo Lopes, Raphael and Karagol Ayan, Burcu and Salimans, Tim and others},
  journal={Advances in Neural Information Processing Systems},
  volume={35},
  pages={36479--36494},
  year={2022}
}

@inproceedings{rombach2022high,
  title={High-resolution image synthesis with latent diffusion models},
  author={Rombach, Robin and Blattmann, Andreas and Lorenz, Dominik and Esser, Patrick and Ommer, Bj{\"o}rn},
  booktitle={Proceedings of the IEEE/CVF Conference on Computer Vision and Pattern Recognition},
  pages={10684--10695},
  year={2022}
}

@article{beutel2020flower,
  title={Flower: A Friendly Federated Learning Research Framework},
  author={Beutel, Daniel J and Topal, Taner and Mathur, Akhil and Qiu, Xinchi and Fernandez-Marques, Javier and Gao, Yan and Sani, Lorenzo and Kwing, Hei Li and Parcollet, Titouan and Gusmão, Pedro PB de and Lane, Nicholas D}, 
  journal={arXiv preprint arXiv:2007.14390},
  year={2020}
}

@misc{von-platen-etal-2022-diffusers,
  author = {Patrick von Platen and Suraj Patil and Anton Lozhkov and Pedro Cuenca and Nathan Lambert and Kashif Rasul and Mishig Davaadorj and Thomas Wolf},
  title = {Diffusers: State-of-the-art diffusion models},
  year = {2022},
  publisher = {GitHub},
  journal = {GitHub repository},
  howpublished = {\url{https://github.com/huggingface/diffusers}}
}

@inproceedings{NIPS2017_8a1d6947,
 author = {Heusel, Martin and Ramsauer, Hubert and Unterthiner, Thomas and Nessler, Bernhard and Hochreiter, Sepp},
 booktitle = {Advances in Neural Information Processing Systems},
 editor = {I. Guyon and U. Von Luxburg and S. Bengio and H. Wallach and R. Fergus and S. Vishwanathan and R. Garnett},
 pages = {},
 publisher = {Curran Associates, Inc.},
 title = {GANs Trained by a Two Time-Scale Update Rule Converge to a Local Nash Equilibrium},
 url = {https://proceedings.neurips.cc/paper_files/paper/2017/file/8a1d694707eb0fefe65871369074926d-Paper.pdf},
 volume = {30},
 year = {2017}
}

@software{Nicki_Skafte_Detlefsen_and_Jiri_Borovec_and_Justus_Schock_and_Ananya_Harsh_and_Teddy_Koker_and_Luca_Di_Liello_and_Daniel_Stancl_and_Changsheng_Quan_and_Maxim_Grechkin_and_William_Falcon_TorchMetrics_-_Measuring_2022,
author = {{Nicki Skafte Detlefsen} and {Jiri Borovec} and {Justus Schock} and {Ananya Harsh} and {Teddy Koker} and {Luca Di Liello} and {Daniel Stancl} and {Changsheng Quan} and {Maxim Grechkin} and {William Falcon}},
doi = {10.21105/joss.04101},
license = {Apache-2.0},
month = feb,
title = {{TorchMetrics - Measuring Reproducibility in PyTorch}},
url = {https://github.com/Lightning-AI/metrics},
year = {2022}
}

@misc{wandb,
title = {Experiment Tracking with Weights and Biases},
year = {2020},
note = {Software available from wandb.com},
url={https://www.wandb.com/},
author = {Biewald, Lukas},
}

@article{kazerouni2022diffusion,
  title={Diffusion models for medical image analysis: A comprehensive survey},
  author={Kazerouni, Amirhossein and Aghdam, Ehsan Khodapanah and Heidari, Moein and Azad, Reza and Fayyaz, Mohsen and Hacihaliloglu, Ilker and Merhof, Dorit},
  journal={arXiv preprint arXiv:2211.07804},
  year={2022}
}

@article{akrout2023diffusion,
  title={Diffusion-based Data Augmentation for Skin Disease Classification: Impact Across Original Medical Datasets to Fully Synthetic Images},
  author={Akrout, Mohamed and Gyepesi, B{\'a}lint and Holl{\'o}, P{\'e}ter and Po{\'o}r, Adrienn and Kincs{\H{o}}, Bl{\'a}ga and Solis, Stephen and Cirone, Katrina and Kawahara, Jeremy and Slade, Dekker and Abid, Latif and others},
  journal={arXiv preprint arXiv:2301.04802},
  year={2023}
}

@article{goodfellow2020generative,
  title={Generative adversarial networks},
  author={Goodfellow, Ian and Pouget-Abadie, Jean and Mirza, Mehdi and Xu, Bing and Warde-Farley, David and Ozair, Sherjil and Courville, Aaron and Bengio, Yoshua},
  journal={Communications of the ACM},
  volume={63},
  number={11},
  pages={139--144},
  year={2020},
  publisher={ACM New York, NY, USA}
}

@inproceedings{rezende2015variational,
  title={Variational inference with normalizing flows},
  author={Rezende, Danilo and Mohamed, Shakir},
  booktitle={International conference on machine learning},
  pages={1530--1538},
  year={2015},
  organization={PMLR}
}

@article{Hou:23,
author = {Benjamin Hou},
journal = {Biomed. Opt. Express},
keywords = {Computed tomography; Image processing; Image quality; Medical imaging; Retina scanning; X ray imaging},
number = {2},
pages = {533--549},
publisher = {Optica Publishing Group},
title = {High-fidelity diabetic retina fundus image synthesis from freestyle lesion maps},
volume = {14},
month = {Feb},
year = {2023},
url = {https://opg.optica.org/boe/abstract.cfm?URI=boe-14-2-533},
doi = {10.1364/BOE.477906},
}

@inproceedings{moghadam2023morphology,
  title={A morphology focused diffusion probabilistic model for synthesis of histopathology images},
  author={Moghadam, Puria Azadi and Van Dalen, Sanne and Martin, Karina C and Lennerz, Jochen and Yip, Stephen and Farahani, Hossein and Bashashati, Ali},
  booktitle={Proceedings of the IEEE/CVF Winter Conference on Applications of Computer Vision},
  pages={2000--2009},
  year={2023}
}

@inproceedings{horvath2022metgan,
  title={METGAN: generative tumour inpainting and modality synthesis in light sheet microscopy},
  author={Horvath, Izabela and Paetzold, Johannes and Schoppe, Oliver and Al-Maskari, Rami and Ezhov, Ivan and Shit, Suprosanna and Li, Hongwei and Ert{\"u}rk, Ali and Menze, Bjoern},
  booktitle={Proceedings of the IEEE/CVF Winter Conference on Applications of Computer Vision},
  pages={227--237},
  year={2022}
}

@inproceedings{fang2022robust,
  title={Robust federated learning with noisy and heterogeneous clients},
  author={Fang, Xiuwen and Ye, Mang},
  booktitle={Proceedings of the IEEE/CVF Conference on Computer Vision and Pattern Recognition},
  pages={10072--10081},
  year={2022}
}

@inproceedings{gao2022feddc,
  title={Feddc: Federated learning with non-iid data via local drift decoupling and correction},
  author={Gao, Liang and Fu, Huazhu and Li, Li and Chen, Yingwen and Xu, Ming and Xu, Cheng-Zhong},
  booktitle={Proceedings of the IEEE/CVF Conference on Computer Vision and Pattern Recognition},
  pages={10112--10121},
  year={2022}
}

@inproceedings{huang2022learn,
  title={Learn from others and be yourself in heterogeneous federated learning},
  author={Huang, Wenke and Ye, Mang and Du, Bo},
  booktitle={Proceedings of the IEEE/CVF Conference on Computer Vision and Pattern Recognition},
  pages={10143--10153},
  year={2022}
}

@inproceedings{jain2023federated,
  title={Federated Learning for Commercial Image Sources},
  author={Jain, Shreyansh and Jerripothula, Koteswar Rao},
  booktitle={Proceedings of the IEEE/CVF Winter Conference on Applications of Computer Vision},
  pages={6534--6543},
  year={2023}
}

@inproceedings{li2022auditing,
  title={Auditing privacy defenses in federated learning via generative gradient leakage},
  author={Li, Zhuohang and Zhang, Jiaxin and Liu, Luyang and Liu, Jian},
  booktitle={Proceedings of the IEEE/CVF Conference on Computer Vision and Pattern Recognition},
  pages={10132--10142},
  year={2022}
}

@inproceedings{li2022federated,
  title={Federated learning with position-aware neurons},
  author={Li, Xin-Chun and Xu, Yi-Chu and Song, Shaoming and Li, Bingshuai and Li, Yinchuan and Shao, Yunfeng and Zhan, De-Chuan},
  booktitle={Proceedings of the IEEE/CVF Conference on Computer Vision and Pattern Recognition},
  pages={10082--10091},
  year={2022}
}

@inproceedings{mendieta2022local,
  title={Local learning matters: Rethinking data heterogeneity in federated learning},
  author={Mendieta, Matias and Yang, Taojiannan and Wang, Pu and Lee, Minwoo and Ding, Zhengming and Chen, Chen},
  booktitle={Proceedings of the IEEE/CVF Conference on Computer Vision and Pattern Recognition},
  pages={8397--8406},
  year={2022}
}

@inproceedings{michieli2021all,
  title={Are all users treated fairly in federated learning systems?},
  author={Michieli, Umberto and Ozay, Mete},
  booktitle={Proceedings of the IEEE/CVF Conference on Computer Vision and Pattern Recognition},
  pages={2318--2322},
  year={2021}
}

@inproceedings{qu2022rethinking,
  title={Rethinking architecture design for tackling data heterogeneity in federated learning},
  author={Qu, Liangqiong and Zhou, Yuyin and Liang, Paul Pu and Xia, Yingda and Wang, Feifei and Adeli, Ehsan and Fei-Fei, Li and Rubin, Daniel},
  booktitle={Proceedings of the IEEE/CVF Conference on Computer Vision and Pattern Recognition},
  pages={10061--10071},
  year={2022}
}

@inproceedings{zhang2021federated,
  title={Federated learning for non-iid data via unified feature learning and optimization objective alignment},
  author={Zhang, Lin and Luo, Yong and Bai, Yan and Du, Bo and Duan, Ling-Yu},
  booktitle={Proceedings of the IEEE/CVF international conference on computer vision},
  pages={4420--4428},
  year={2021}
}

@article{hatamizadeh2023gradient,
  title={Do gradient inversion attacks make federated learning unsafe?},
  author={Hatamizadeh, Ali and Yin, Hongxu and Molchanov, Pavlo and Myronenko, Andriy and Li, Wenqi and Dogra, Prerna and Feng, Andrew and Flores, Mona G and Kautz, Jan and Xu, Daguang and others},
  journal={IEEE Transactions on Medical Imaging},
  year={2023},
  publisher={IEEE}
}

@article{hosseini2023proportionally,
  title={Proportionally Fair Hospital Collaborations in Federated Learning of Histopathology Images},
  author={Hosseini, S Maryam and Sikaroudi, Milad and Babaie, Morteza and Tizhoosh, HR},
  journal={IEEE Transactions on Medical Imaging},
  year={2023},
  publisher={IEEE}
}

@article{nguyen2022federated,
  title={Federated learning for smart healthcare: A survey},
  author={Nguyen, Dinh C and Pham, Quoc-Viet and Pathirana, Pubudu N and Ding, Ming and Seneviratne, Aruna and Lin, Zihuai and Dobre, Octavia and Hwang, Won-Joo},
  journal={ACM Computing Surveys (CSUR)},
  volume={55},
  number={3},
  pages={1--37},
  year={2022},
  publisher={ACM New York, NY}
}

@inproceedings{li2021model,
  title={Model-contrastive federated learning},
  author={Li, Qinbin and He, Bingsheng and Song, Dawn},
  booktitle={Proceedings of the IEEE/CVF Conference on Computer Vision and Pattern Recognition},
  pages={10713--10722},
  year={2021}
}

@article{rieke2020future,
  title={The future of digital health with federated learning},
  author={Rieke, Nicola and Hancox, Jonny and Li, Wenqi and Milletari, Fausto and Roth, Holger R and Albarqouni, Shadi and Bakas, Spyridon and Galtier, Mathieu N and Landman, Bennett A and Maier-Hein, Klaus and others},
  journal={NPJ digital medicine},
  volume={3},
  number={1},
  pages={119},
  year={2020},
  publisher={Nature Publishing Group UK London}
}

@inproceedings{qin2023class,
  title={Class-Balancing Diffusion Models},
  author={Qin, Yiming and Zheng, Huangjie and Yao, Jiangchao and Zhou, Mingyuan and Zhang, Ya},
  booktitle={Proceedings of the IEEE/CVF Conference on Computer Vision and Pattern Recognition},
  pages={18434--18443},
  year={2023}
}

@inproceedings{hu2023self,
  title={Self-guided diffusion models},
  author={Hu, Vincent Tao and Zhang, David W and Asano, Yuki M and Burghouts, Gertjan J and Snoek, Cees GM},
  booktitle={Proceedings of the IEEE/CVF Conference on Computer Vision and Pattern Recognition},
  pages={18413--18422},
  year={2023}
}
