## Supplementary figures and images for "FMed-Diffusion Federated Learning on Medical Image Diffusion"

### C1TrainLoss.jpg

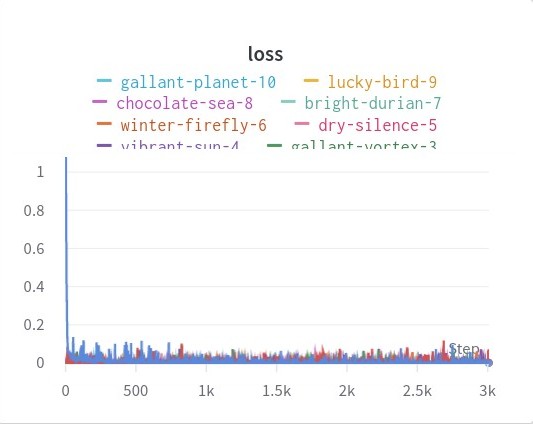

### C2TrainLoss.jpg

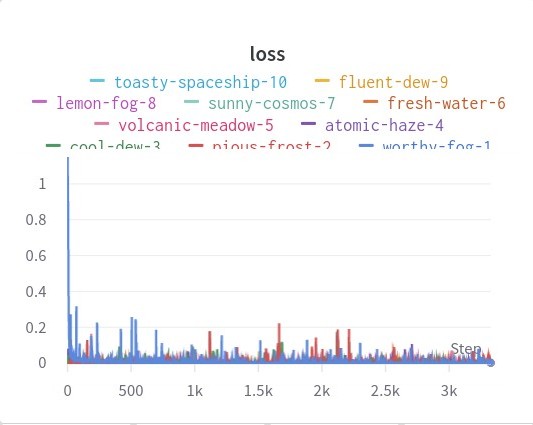

### C3TrainLoss.jpg

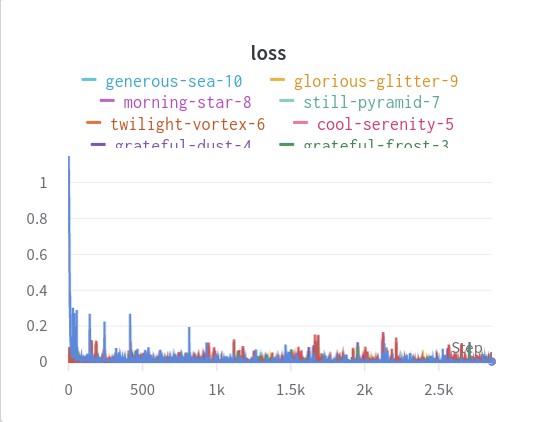

### C_s12.png

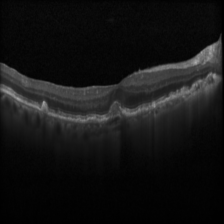

### C_s16.png

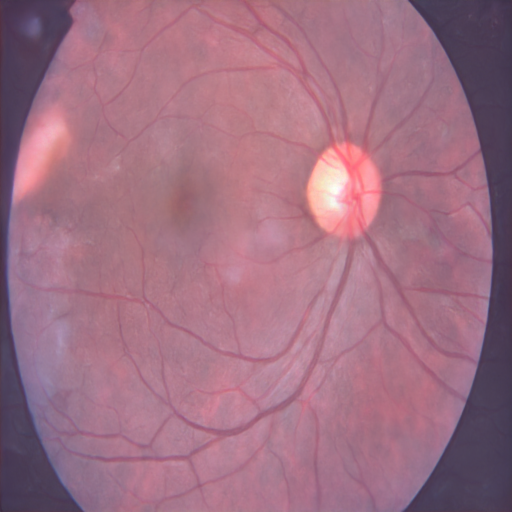

### C_s16_COVID.png

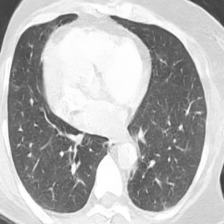

### CT24.png

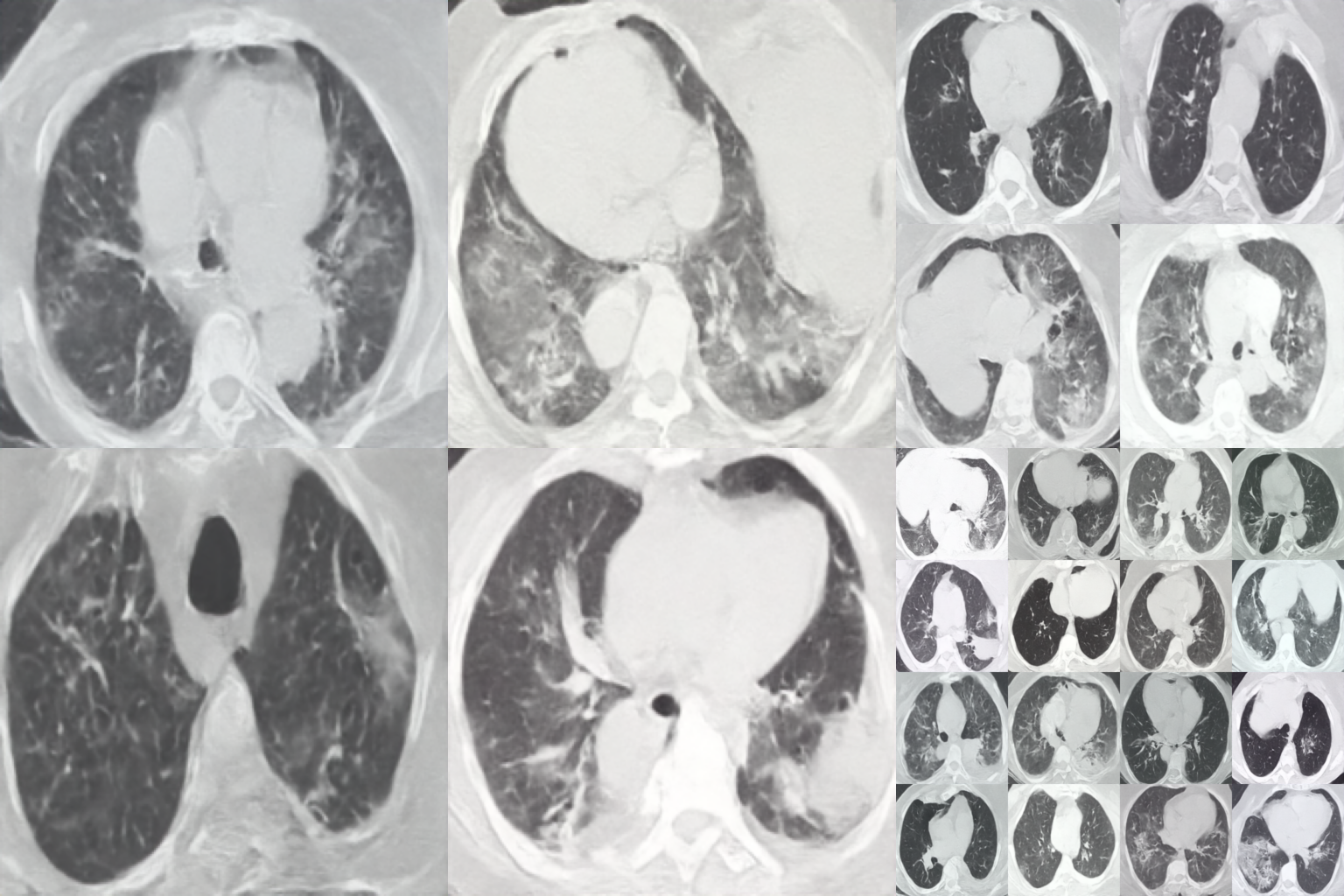

### Diff.pdf

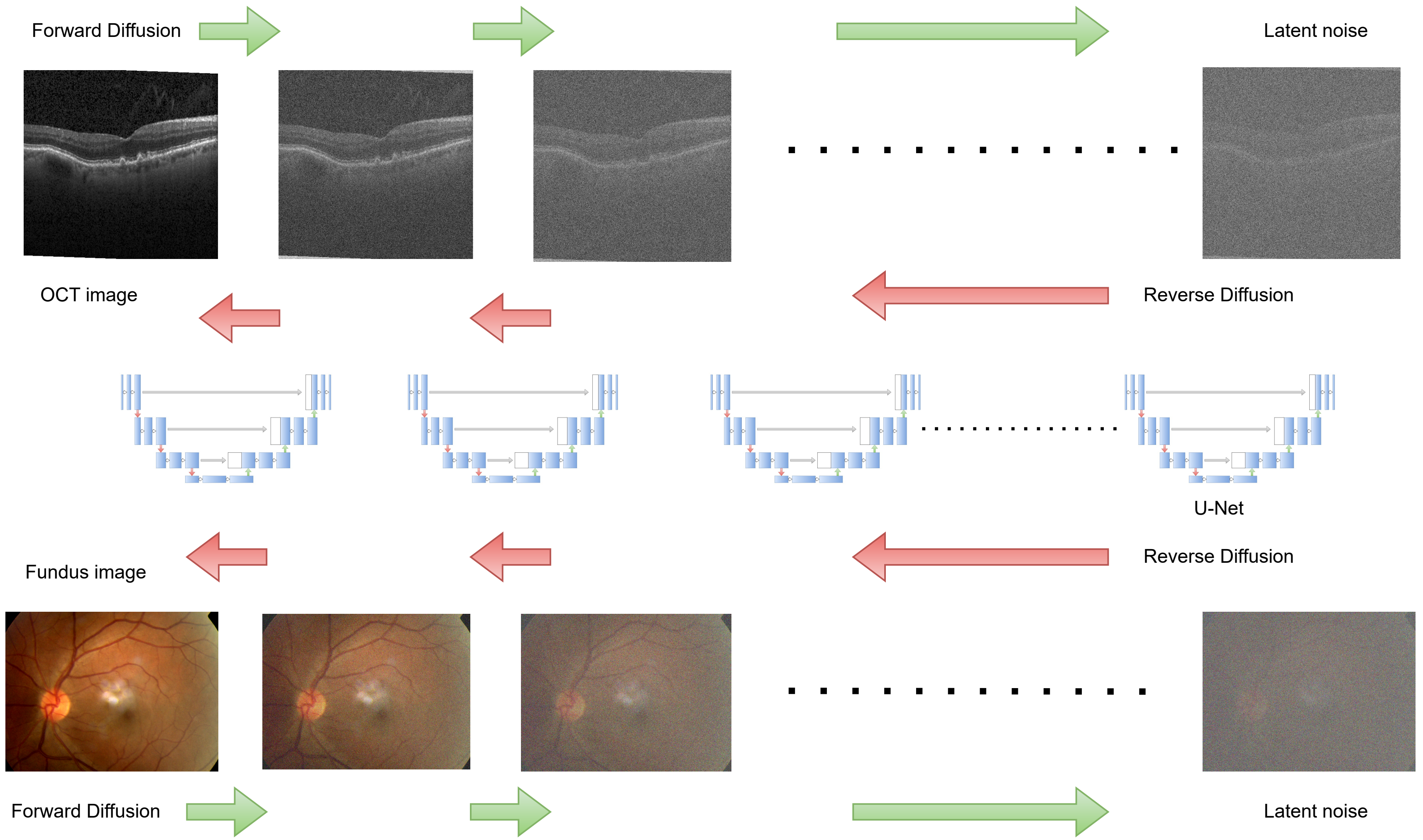

### Dr24.png

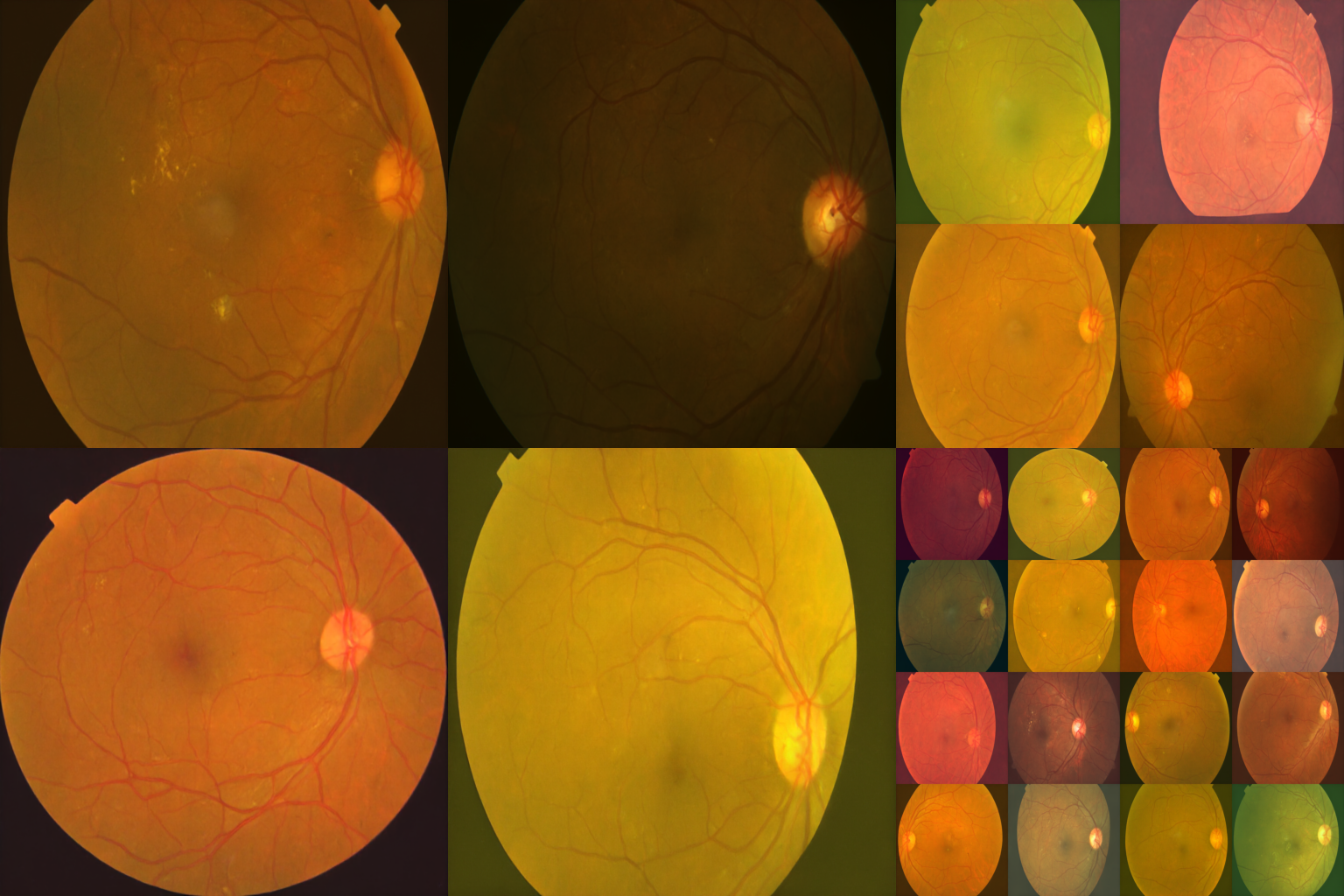

### F_s11.png

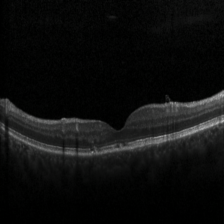

### F_s46.png

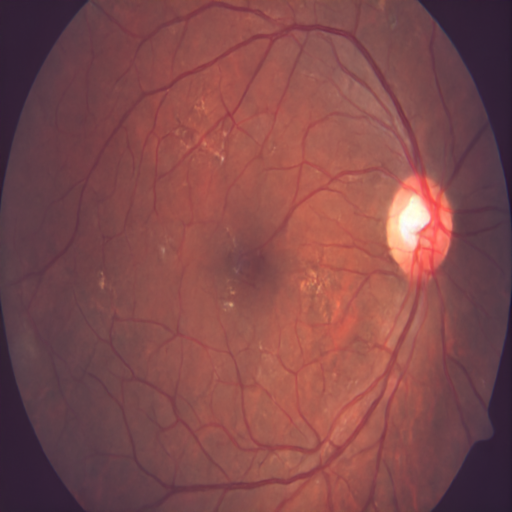

### F_s187.png

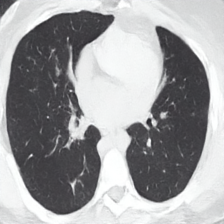

### FL1.drawio.pdf

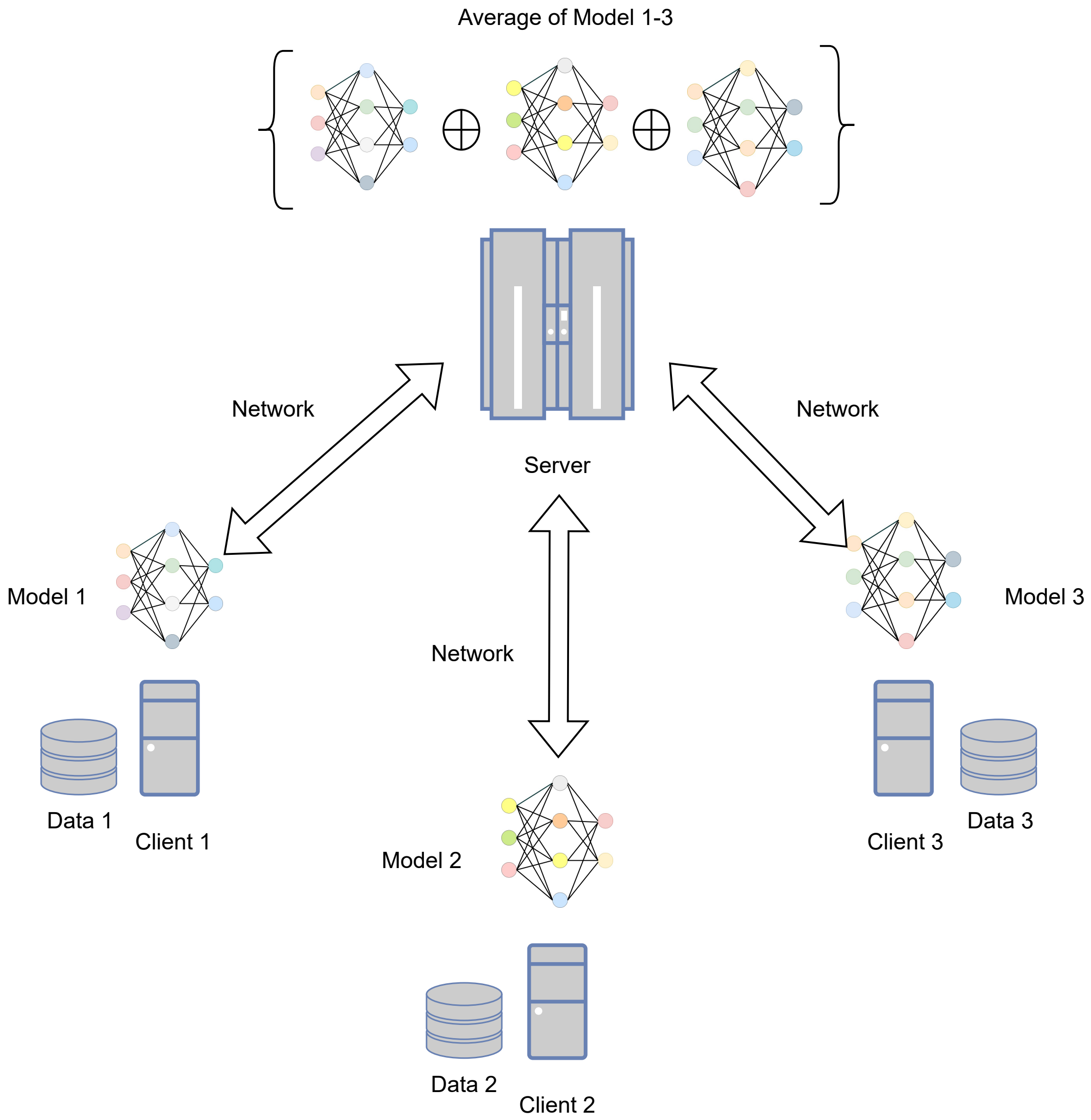

### line_graph_FID.png

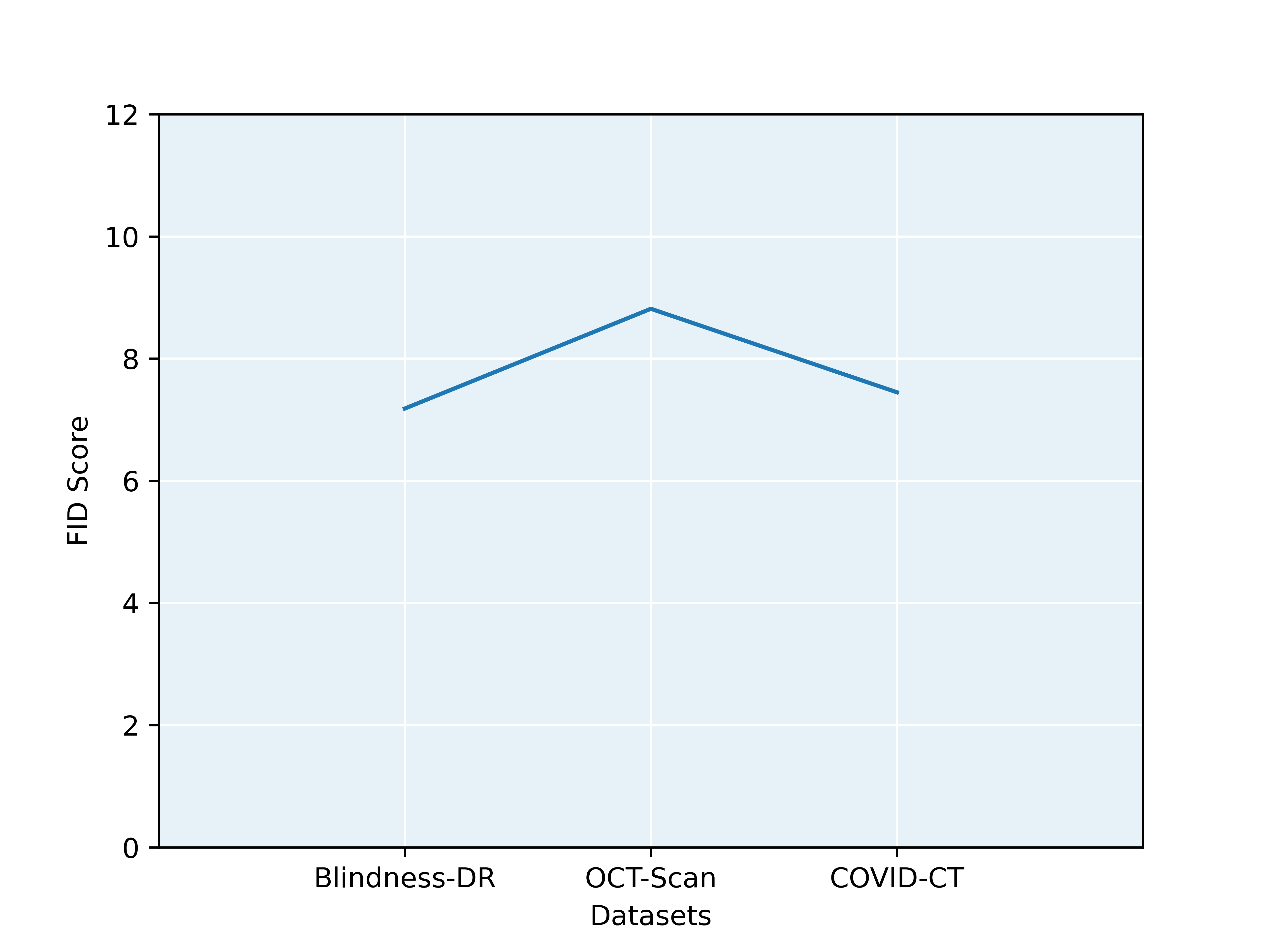

### muru.jpg

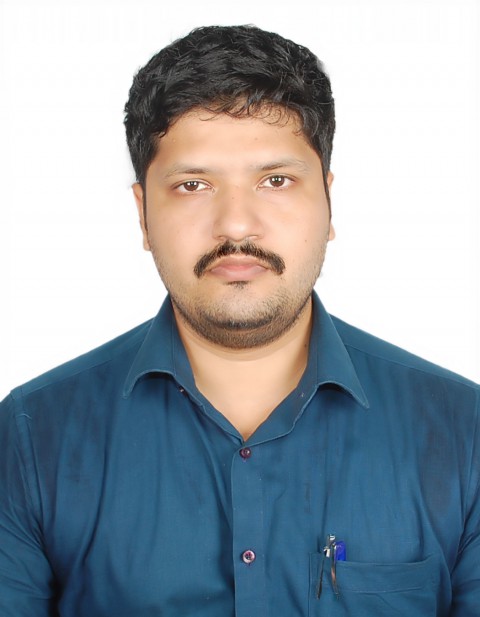

### OCT24.png

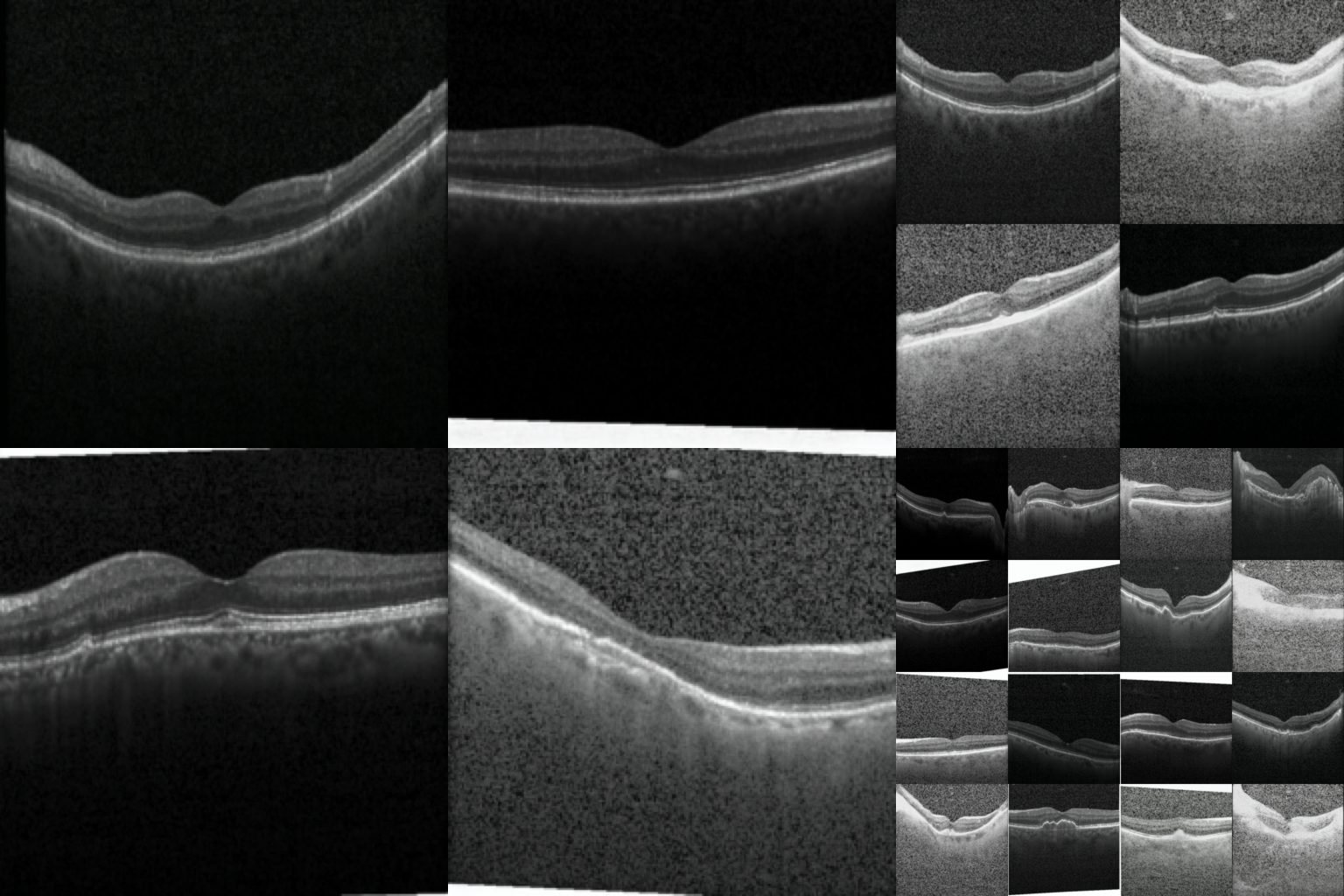
